# A computational understanding of zoomorphic perception in the human brain

**DOI:** 10.1101/2022.09.26.509447

**Authors:** Stefanie Duyck, Stefania Bracci, Hans Op de Beeck

## Abstract

It is common to find objects that resemble animals on purpose (e.g., toys). While the perception of such objects as animal-like seems obvious to humans, such “Animal bias” for zoomorphic objects turned out to be a striking discrepancy between the human brain and artificial visual systems known as deep neural networks (DNNs). We provide a computational understanding of the human Animal bias. We successfully induced this bias in DNNs trained explicitly with zoomorphic objects. Alternative training schedules, focusing on previously identified differences between the brain and DNNs, failed to cause an Animal bias. Specifically, we considered the superordinate distinction between animate and inanimate classes, the sensitivity for faces and bodies, the bias for shape over texture, and the role of ecologically valid categories. These findings provide computational support that the Animal bias for zoomorphic objects is a unique property of human perception yet can be explained by human learning history.

## Introduction

The life of babies and children is full of zoomorphic objects: objects made to look like animals, such as a cuddly toy or a butterfly-shaped rattle. This practice might be as old as human culture itself, judging from archeological research that has revealed plenty of prehistoric artefacts with animal characteristics, e.g., pottery vessels and jewelry. In addition, the human mind does not need many cues to interpret even simple visual shapes as animate, such as when two triangles chase a square (1–4) Previous research has suggested various benefits^5,6^ and origins of this animacy perception (5) (6). However, despite the prevalence and importance of zoomorphism, we lack a computational understanding of this perceptual phenomenon: How and why does the perception of animacy in zoomorphic objects come about?

Previous research has provided insights into how the distinction between animals and other object categories is represented in the human brain. In occipitotemporal cortex (VTC) there are distinct neural representations of categories such as faces (7,8), bodies (9,10), and animals (11,12). At the broader level, studies have found a hierarchically organized animacy continuum in VTC (13–16), which has been related to perceptual and/or conceptual properties of animals (15–19).

Recent research revealed that the responses in VTC to zoomorphic objects explain why humans are so prone to interpret these objects as animate (17). Representational similarity analyses showed that anterior VTC represents zoomorphic objects more like animals than like regular objects, which we here refer to as an “Animal bias”. The effect is reminiscent of other findings with face pareidolia, the tendency to see faces in objects: VTC tends to represent objects that trigger face pareidolia a bit more like faces than other objects (20). However, the bias was much more prominent with zoomorphic objects (17): Zoomorphic objects were represented as animals, not somewhere in between animals and regular objects.

These neuroscientific investigations also exposed the extent to which we lack a computational understanding of why the human visual system is so prone to this Animal bias. Many recent studies have suggested that DNNs trained on image classification provide a useful computational model of human vision (17,21–26). More specifically, the later layers (i.e., fully connected or deep layers) of these networks typically show a similar representational similarity structure as observed in VTC (27), including a rudimentary animacy continuum representation (16,21,22). However, there is a striking discrepancy between human and DNN vision when it comes to zoomorphic objects. DNNs show the exact opposite bias from human visual cortex, and represent zoomorphic objects as inanimate, even as inanimate as regular objects, a so-called Object bias (17). Thus, while DNNs might be a useful computational model of human perception in general, they totally fail as a model for zoomorphic perception.

Here we investigate what it takes to construct a computational model that represents zoomorphic objects in a similar manner as the human visual system, with an Animal bias rather than an Object bias. We test several hypotheses. First, we investigate whether it is possible to induce an Animal bias in DNNs by explicitly training DNNs to categorize zoomorphic objects as animals, contrary to their initial Object bias. This effort was successful. This success suggests a first hypothesis for why the adult human brain might show a strong Animal bias: because of the frequent encounter of zoomorphic objects meant to represent animals.

However, there are many other properties of the human visual system and human visual experience that might more indirectly induce an Animal bias. In comparison to DNNs, human vision has been shown to be particularly sensitive to the superordinate distinction between animals and other objects (13–16), to categories like faces and bodies (7–10), to global shape (28), and to the natural statistics in the environment (29). None of these properties is captured fully in the benchmark DNN models in the literature. We tested computational models constructed to remediate these shortcomings, but none of them displayed an Animal bias in DNNs, nor captured brain representations well. Through these computational tests we narrow down the possible causes of the Animal bias to the relevance of zoomorphic objects during past training of the human visual system.

## Results

### Fine-tuning shifts the representational space for zoomorphic objects in pretrained DNNs

To shift the representational space for zoomorphic objects, pretrained AlexNet networks (PT ImageNet) were fine-tuned (FT) over the last two fully connected layers (i.e., FC7 and FC8). A dataset with three separate categories (i.e., animals, zoomorphic objects and objects) was split for training, validating and testing the networks (Fig. 1a). Depending on the type of training (Fig. 1b), two out of three categories were joined together. The FT Animal bias DNN was trained to classify images either as animals&zoomorphic or as objects. Conversely, FT Object bias DNN, was trained on animals versus zoomorphic&objects. The test set contained images from earlier work (17) (Fig. 1c) and new ones (for a total of 99 test images, 33 per category). Analyses on DNNs data are performed on the entire test set, whereas analyses investigating the correlations with the neural data are performed on the subset derived from earlier work. All results are based upon the normalized distance ((1 – Pearson R)/mean RDM) between the output representations of all image pairs, resulting in representational dissimilarity matrices (RDMs) (Fig. 2a). All p-values smaller than .01 are considered significant.

**Fig 1.**
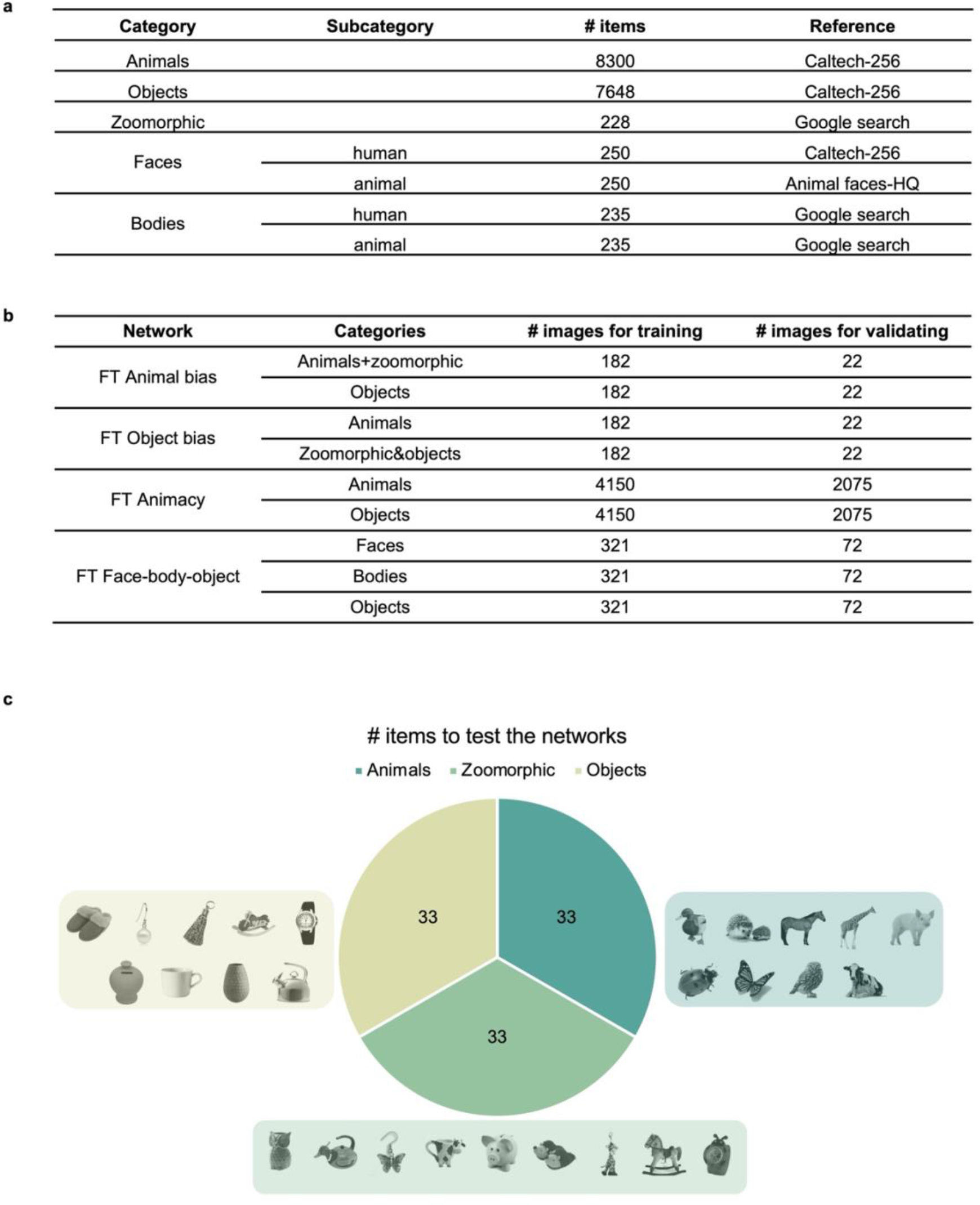
Overview of the stimuli. **a** Overview of categories in each type of training regime and the number of images used for training and validating. **b** Info on the whereabouts and numbers of all image categories. **c** An overview on the number of images in the test phase, used to construct the RDMs. 33 images in each category, of which the 9 Bracci subset images are displayed for each category.

**Fig. 2.**
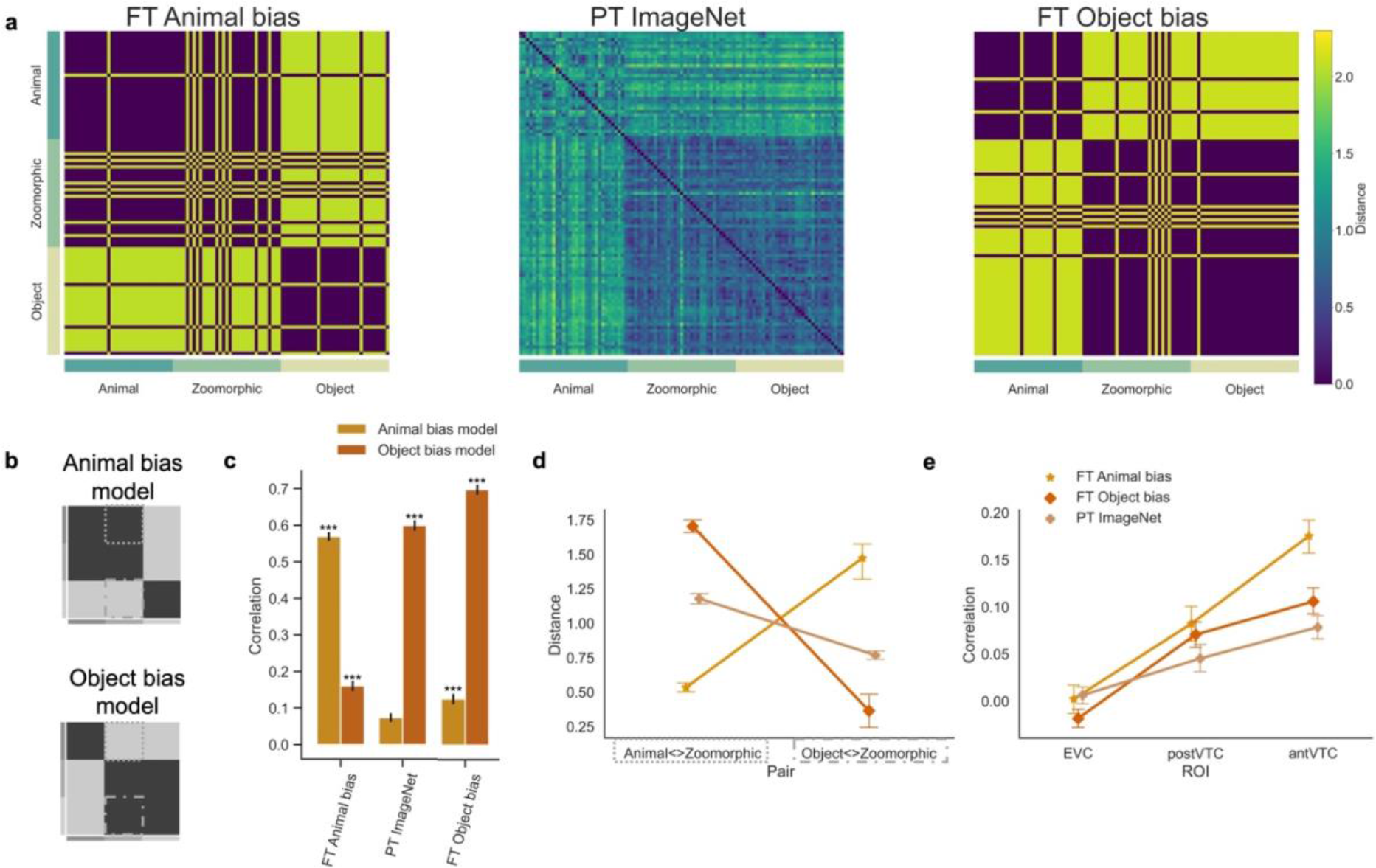
Overview of the explicit Animal bias training results. **a** The RDMs in FC8 of three DNNs included in the analyses are displayed. FT = fine-tuned AlexNet, PT = pretrained AlexNet. For FC8 in these networks the RDMs are binary because there are only 2 output units. **b** Graphical illustration of the pre-defined conceptual Animal and Object bias model. **c** The graph represents the correlation for each DNN with each bias model. Significant values (i.e., *** p < .0001, ** p < .001, * p < .01), were computed with permutation tests (10,000 randomizations), and error bars indicate standard error computed via bootstrapping. **d** Mean distance score of each image in the contrast Animal<>Zoomorphic and Object<>Zoomorphic (see dotted outlines in panel b) and this for each DNN. **e** Graph with the correlations between the individual neural data in the three regions of interest (i.e., EVC, posterior- and anterior VTC) and the DNN data resulting from the different training regimes. All error bars represent the standard error.

These matrices showed a strong effect of the fine-tuning training protocol in terms of correlations with pre-specified conceptual representational structures (see Fig. 2b), which aligned either with an Object bias (Object bias model) or with the human-like Animal bias (Animal bias model). More specifically, (Fig. 2c) the object space derived from the FT Animal bias DNNs was strongly correlated with the Animal bias model (r = 0.57) relative to the Object bias model (r = 0.16; p < .001). The opposite trend was observed for the FT Object bias DNN, whose object space was strongly correlated with the Object bias model (r = 0.70) relative to the Animal bias model (r = 0.13; p < .001). The representational space derived from the baseline PT ImageNet network was also correlated more strongly with the Object bias model (r = .60; Animal bias model: r = .07, p < .001). We restrict the reporting throughout this Results section to one network architecture (AlexNet), and the last network layer (FC8 in AlexNet), but similar trends (though with smaller effect size) are present in layer FC7 (Supplementary Fig. 2), and similar conclusions are supported by simulations with another frequently used architecture (VGG, Supplementary Fig. 1).

In addition to correlations with the independent models, that include all non-diagonal cells in the RDM, we specifically investigated (Fig. 2d) the distance between animal and zoomorphic objects, and between regular and zoomorphic objects, both averaged across all pair-wise image comparisons (see dotted outlines in Fig. 2b). The smaller the distance, the more similar the representations of both categories are. An animal bias would manifest itself by a smaller distance in Animal<>Zoomorphic than in Zoomorphic<>object. A factorial repeated-measures ANOVA on the dissimilarity scores with factors Distances (Animal<>Zoomorphic; Object<>Zoomorphic) x Network (PT ImageNet; FT Object bias; FT Animal bias) revealed a significant interaction effect (F(2,64) = 94.88, p < .001). Post-hoc analyses further investigated the degree to which the different networks learnt to consider zoomorphic objects more like animals or objects. Relative to the pretrained PT ImageNet DNN, the finetuning in the FT Animal bias DNN moved the zoomorphic objects closer to the animals (M = 0.53, t(32) = 20.07, p_corr_ < .001) and further away from objects (M = 1.47, t(32) = -5.15, p_corr_ < .001). The reverse is observed for the FT Object bias DNN, with zoomorphic farther from animals (M = 1.70, t(32) = -6.06, p_corr_ < .001) and closer to objects (M = 0.37, t(32) = 4.26, p_corr_ < .001) compared to PT ImageNet. In sum, the fine-tuned networks shifted towards the bias they were trained in, with particularly large effect size for the Animal bias training.

### Animal bias DNN explains brain representations better than Object bias DNNs

The earlier finding (17) of a strong Animal bias in anterior VTC, makes us predict that the FT Animal bias DNN should correlate well with the representational similarity in ant VTC. The individual RDMs (n = 16) of each region of interest (early visual cortex (EVC), anterior and posterior VTC) were correlated with the DNN RDMs (Fig. 2e). In VTC, in contrast to EVC, all DNNs showed a positive correlation (one sided one sample t-test, all p <= 0.003) with the neural data in anterior and posterior VTC. This result converges with previous results showing that fully connected layers in DNNs better predict brain representations in high level visual areas relative to low-level visual cortex (30). Importantly, we observed by means of a repeated-measures ANOVA, a significant interaction between ROI (EVC; ant VTC; post VTC) x Network (FT Animal bias; PT ImageNet; FT Object bias) (F(2,60) = 7.93, p = .002). Post-hoc analyses provided insight in representations learned by the FT Animal bias DNNs, such that these were significantly more correlated with representations in ant VTC (r = 0.18), relative to representations learned by FT Object bias (r = 0.11, t(15) = 3.86, p_corr_ = .007) and the standard PT ImageNet DNN (r = 0.08, t(15) = 5.34, p_corr_ < .001). This difference was not observed in post VTC (Object bias: r = 0.07, t(15) = 0.89, p_corr_ = 0.43; PT ImageNet: r = 0.05, t(15) = 2.47, p_corr_ = .08). Together these results show that a DNN trained to classify zoomorphic objects like animals can capture the human anterior VTC representations much better than DNNs without such training.

Overall, the findings obtained after training a network to classify zoomorphic objects as animals, showed that DNNs can show an Animal bias and resulting representational structure mimics brain representations to a greater extent relative to benchmark DNNs. However, the failure of benchmark DNNs to show this Animal bias might also have resulted from other obvious and previously described differences in the visual input and tasks that humans and artificial systems are exposed to. These differences are more generic than what happens with zoomorphic objects, but they might indirectly cause a particular processing of such objects. In the following sections, we test several proposals. In each case, we compare an additional DNN trained to test the hypothesis at hand with the baseline PT ImageNet DNN and the FT Animal bias DNN. As such, all ANOVA analyses with the factor ‘Network’ consist of three levels: FT Animal bias DNN, PT ImageNet DNN and the newly added DNN.

### Hypothesis one: Animal bias because of explicit training in categories such as animacy or faces and bodies?

The human visual system is particularly sensitive for the distinction between inanimate and animate objects, the latter represented as a continuum (13–16), and shows regions with a very strong selectivity for faces and bodies (9,31). However, these two types of selectivity are only partially captured by benchmark DNNs, possibly due to limitations in their training history. First, whereas the animacy continuum is present in DNNs, the categorical distinction seems less pronounced than in the human brain (32). DNNs are trained on very specific labels (e.g., specific species of dogs and birds) and not so much on superordinate categories (e.g., animate). Remediating this can in some training regimes improve the fit between DNN representations and responses in visual cortex (33). Fine-tuning the PT ImageNet in the superordinate animate-inanimate distinction (FT Animacy), without the inclusion of zoomorphic stimuli, might give rise to more Animal bias and convergence with VTC for zoomorphic objects.

Second, whereas face and body selectivity is very prominent in the human visual system, these two categories are not included as explicit classes in the typical ImageNet training protocol. Given that face and body selectivity is an important determinant of animacy representations (16), it might also underlie the Animal bias. We tested whether fine-tuning a DNN with human and animal faces and bodies, in addition to objects, might induce an Animal bias (FT Face-body-object).

Figure 3 displays the results for FT Animacy and FT Face-body-object, together with the DNNs, PT ImageNet and FT Animal bias. The RDMs from the alternative networks are shown in Fig. 3a, where we can already observe that the representational structure in the two new DNNs tends to align with PT ImageNet and that FT Animal bias stands apart as the odd-one-out. Quantitative analyses revealed strong positive correlations for both alternative DNNs with the independent Object bias model (Fig. 3b; FT Animacy DNN: r = .53; FT Face-body-object DNN: r = 0.48) relative to the Animal bias model (all p < .001) which did not differ from zero (FT Animacy DNN: r = .03, p = .10; FT Face-body-object DNN: r = 0.008, p = .34).

**Fig. 3.**
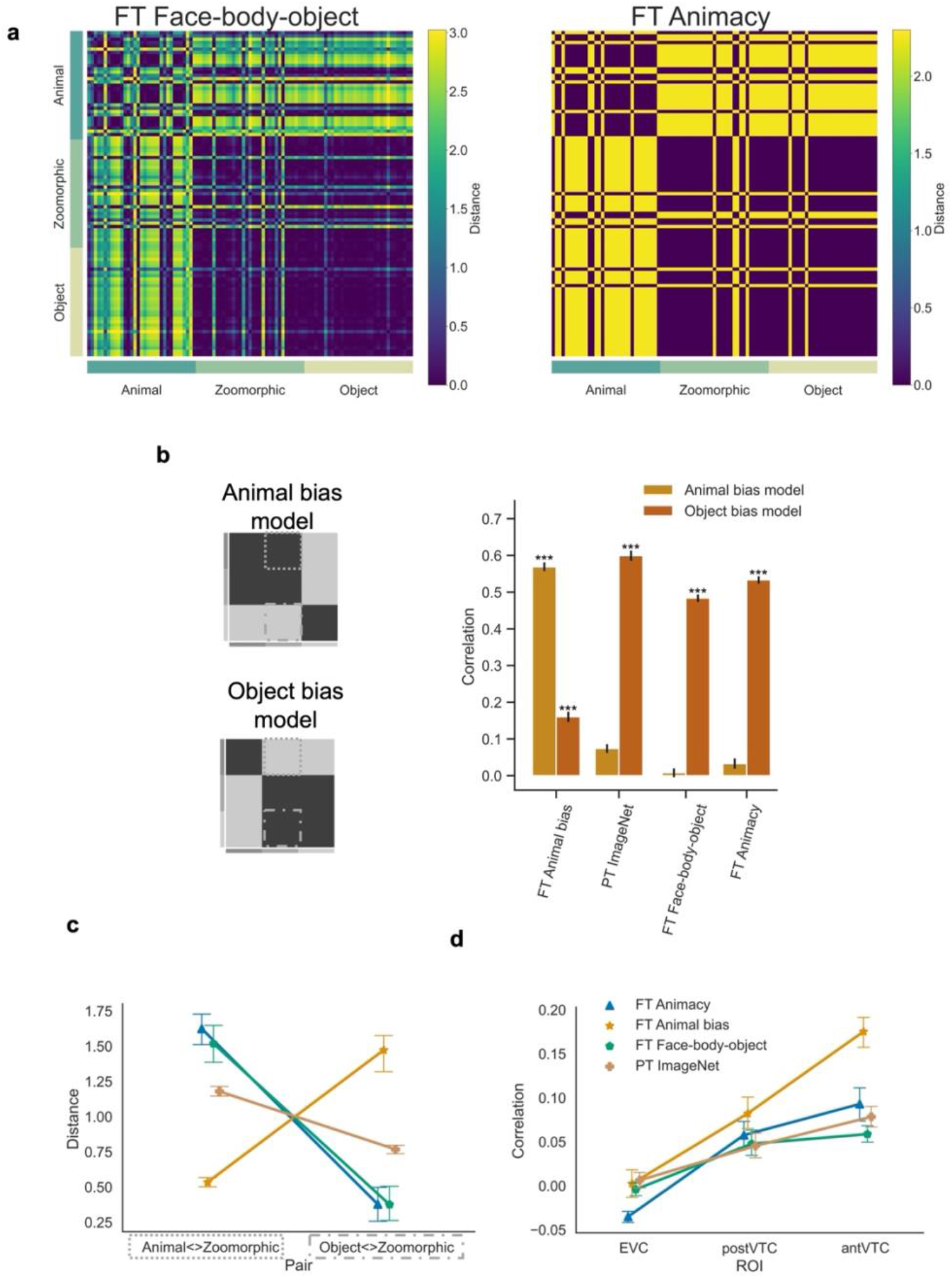
Alternative hypothesis one: sensitivity for specific categories. **a** The RDMs of two alternative networks included in the analyses are displayed. FT = fine-tuned AlexNet, PT = pretrained AlexNet. **b** Graphical illustration of the pre-defined conceptual Animal and Object bias model. The graph represents the correlation for each DNN with each bias model. Significant values (i.e., *** p < .0001, ** p < .001, * p < .01), were computed with permutation tests (10,000 randomizations), and error bars indicate standard error computed via bootstrapping. **c** Mean distance score of each image in the contrast Animal<>Zoomorphic and Object<>Zoomorphic and this for each DNN. **d** Graph with the individual correlations between the neural data in the three regions of interest (i.e., EVC, posterior- and anterior VTC) and the DNN data resulting from the different training regimes. All error bars represent the standard error.

Next, we investigated the distance between zoomorphic objects and either animals or objects (Fig. 3c), as calculated in the previous results section. The ANOVA with Distances (Animal<>Zoomorphic; Object<>Zoomorphic) x Network revealed significant interaction terms for both alternative DNNs (FT Animacy DNN: F(2,64) = 94.88, p < .001; FT Face-body-object DNN: F(2,64) = 73.33, p < .001). Post-hoc analyses showed that relative to the pretrained ImageNet DNN, the distance between animals and zoomorphic objects (M = 1.18) even increased in both FT Animacy DNN (M = 1.62) and FT Face-body-object DNN (M = 1.52), although only in the former it reached statistical significance (t(32) = 3.01, p_corr_ = .006; t(32) = 0.71, p_corr_ = .56, respectively). At the same time, compared to PT ImageNet DNN, the distance between zoomorphic and regular objects (M = 0.77) significantly decreased (FT Animacy DNN: M = 0.38, t(32) = -6.52, p_corr_ < .001; FT Face-body-object DNN: M = 0.38, t(32) = -5.79, p_corr_ < .001). These findings were significantly opposed relative to representations observed in FT Animal bias DNN, such that in both DNNs (FT Animacy and FT Face-body-object) the distance between animals and zoomorphic objects was significantly larger (FT Animacy: M = 1.62, t(32) = 8.68, p_corr_ < .001; FT Face-body-object: M = 1.52, t(32) = 6.01, p_corr_ < .001), whereas zoomorphic images were represented significantly closer to real objects (FT Animacy: M = 0.38, t(32) = -7.17, p_corr_ < .001; FT Face-body-object: M = 0.38, t(32) = -7.50, p_corr_ < .001).

In terms of correlations with the neural representations (Fig. 3d), none of the alternative trainings (Animacy and Face-Body) resulted in a representational structure that could reach performance obtained by the Animal bias DNN. Specifically, for both alternative DNNs, an ANOVA with factors ROI (EVC; posterior VTC; anterior VTC) and Network (FT Animal bias bias; PT ImageNet; FT Animacy or FT Face-body-object) on the correlations revealed a significant interaction between ROI and Network (*FT Animacy*: F(4,60) = 8.05, p = .002; *FT Face-body-object*: F(4,60) = 11.05, p < .001). Most importantly, in each case, the alternative DNN showed significantly lower correlations in ant VTC compared to FT Animal bias (FT animacy: r_Animacy DNN_ = 0.09, r_Animal bias DNN_ = 0.18, t(15) = -4.83, p_corr_ < .001; FT Face-body-object: r_Face-body-object DNN_ = 0.06, r_Animal bias DNN_ = 0.18, t(15) = -6.84, p_corr_ < .001), while correlations were similar to the ones observed for PT ImageNet (FT animacy: r = 0.08, t(15) = 1.43, p =.20; FT Face-body-object: r = 0.08, t(15) = -2.15, p_corr_ = .09). To conclude, fine-tuning the network to selectivity for animacy or for faces and bodies captured less of the anterior VTC representations than the DNN fine-tuned to Animal bias, and did not improve the fit relative to standard pretrained DNN.

### Hypothesis two: Can the Animal bias emerge in DNNs trained to focus on shape over texture?

Geirhos and colleagues(34) found that DNNs tend to rely more upon texture than shape when categorizing objects (texture bias), while human observers show the opposite bias (shape bias). Given that this finding is one of the best documented discrepancies between human and AI vision, we considered the possibility that the Object bias in DNNs might be related to their texture bias, reducing the two seemingly separate phenomena to one and the same underlying tendency. Geirhos and colleagues (34) also showed that the texture bias in DNNs is converted into a shape bias when using a different training regime with a stimulus set in which images are more variable in texture and style, the so-called Stylized ImageNet. We tested to which extent a DNN pretrained on the Stylized ImageNet (PT Stylized ImageNet) would induce an Animal bias.

Findings showed (Fig. 4b) a positive correlation between the PT Stylized ImageNet DNN and the independent Object bias model (r = 0.52) which was significantly higher relative to the correlation observed with the Animal bias model (r = 0.07, p < .001). Analyzing the representational distance (Fig. 4c) for zoomorphic objects relative to the animals and the objects revealed a representational space like the one observed in the baseline pretrained ImageNet DNN. That is, animals and zoomorphic objects in the PT Stylized DNN (M = 1.17, significant interaction effect F(2,64) = 106.85, p < .001) showed a larger distance compared to FT Animal bias (M = 0.53, t(32) = 19.36, p_corr_ < .001) but a similar distance relative to PT ImageNet (M = 1.18, t(32) = -0.04, p_corr_ = .97). Conversely, there was a smaller distance between zoomorphic and real objects in PT Stylized (M = 0.81) compared to FT Animal bias (M = 1.47, t(32) = -4.29, p_corr_ < .001) and equal distance relative to PT ImageNet (M = 0.77, t(32) = -0.52, p_corr_ = .65). These findings illustrated the absence of a strong Animal bias in PT Stylized ImageNet.

**Fig. 4.**
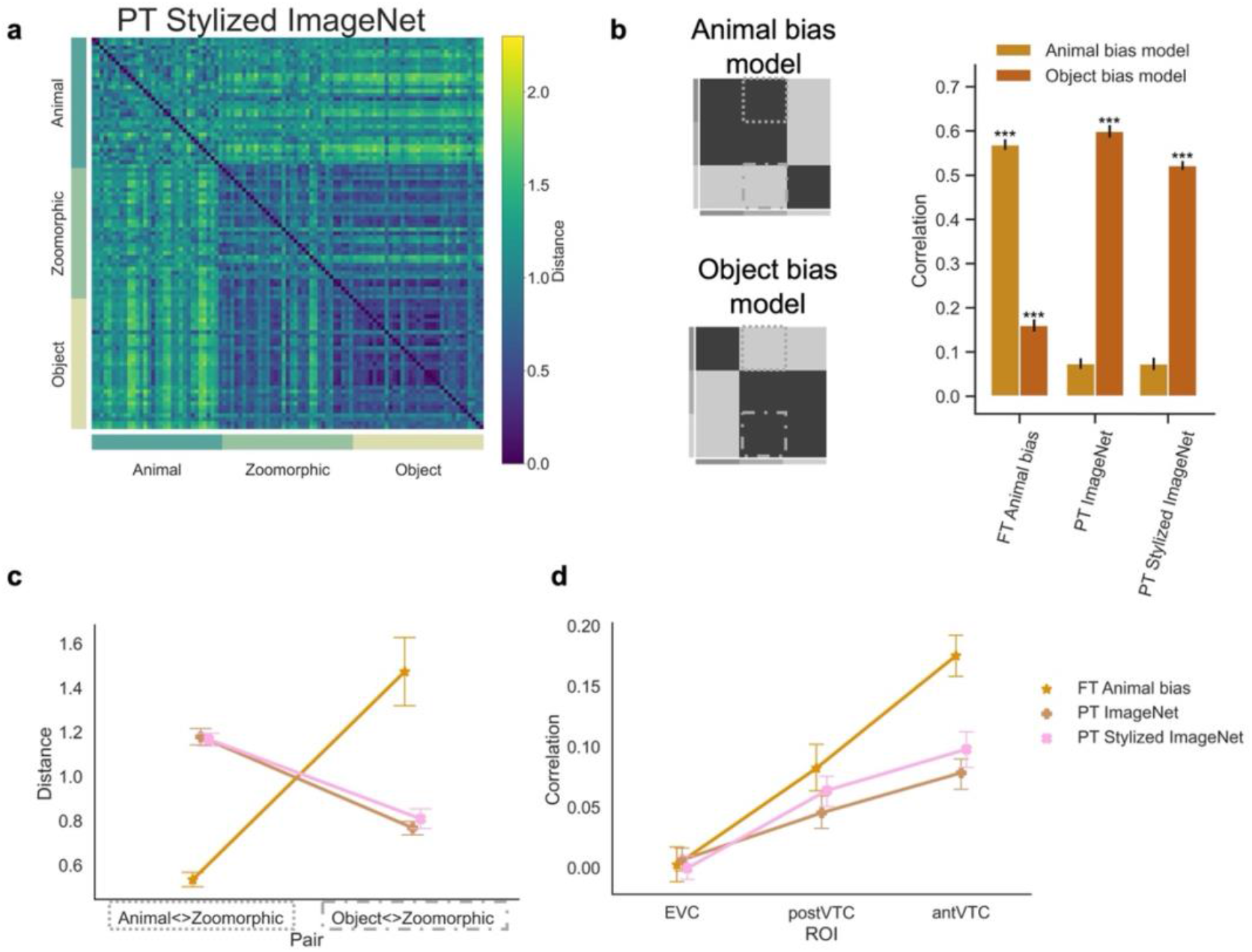
Alternative hypothesis two: tackling shape-texture bias. **a** The pretrained Stylized ImageNet RDM included in the analyses is displayed. PT = pretrained AlexNet, FT = fine-tuned AlexNet. **b** Graphical illustration of the independent Animal and Object bias model. The graph represents the correlation for each DNN with each bias model. Significant values (i.e., *** p < .0001, ** p < .001, * p < .01), were computed with permutation tests (10,000 randomizations), and error bars indicate standard error computed via bootstrapping. **c** Mean distance score of each image in the contrast Animal<>Zoomorphic and Object<>Zoomorphic and this for each DNN. **d** Graph with the individual correlations between the neural data in the three regions of interest (i.e., EVC, posterior- and anterior VTC) and the DNN data resulting from the different training regimes. All error bars represent the standard error.

Next, we looked at the correlation with neural representations (Fig. 4d). A repeated measures ANOVA with factors ROI (EVC; post VTC; ant VTC) x Network (PT Stylized ImageNet; PT ImageNet; FT Animal bias) on the correlation with the neural data revealed significant interacting terms (F(4,60) = 7.99, p = .003). Post-hoc analyses showed that whereas in EVC there was no difference between the three DNNs (all p_corr_ > .36), when moving downstream, this difference increased and reached significance in ant VTC where correlations for the FT Animal bias DNN were significantly higher relative to the PT Stylized DNN (ant VTC: t(15) = 4.54, p_corr_ = .002) and the baseline PT ImageNet DNN (ant VTC: t(15) = 5.34, p_corr_ < .001). In addition, relative to the baseline PT ImageNet DNN, the PT Stylized DNN was slightly better correlated to the neural data in VTC but these differences did not survive correction for multiple comparisons (post VTC; t(15) = 2.71, p_corr_ = .036). To conclude, even a network that is trained to shift to a human-like shape bias does not show a human-like Animal bias.

### Hypothesis three: Animal bias because of network trained with more naturalistic categories?

Finally, we tested the hypothesis that training with more ecologically valid categories would result in a human-like Animal bias. A previous study introduced the ‘Ecoset’ stimulus set and showed that training with Ecoset results in a representational structure in later DNN layers that fits slightly better with brain representations compared to training with ImageNet (35). We tested the effect of this Ecoset training on the representation of zoomorphic objects by testing AlexNet pretrained with Ecoset. Figure 5a displays the RDM obtained for this PT Ecoset DNN, alongside the previously reported RDMs for PT ImageNet and FT Animal bias. The visual similarity between PT Ecoset and PT ImageNet is striking.

**Fig. 5.**
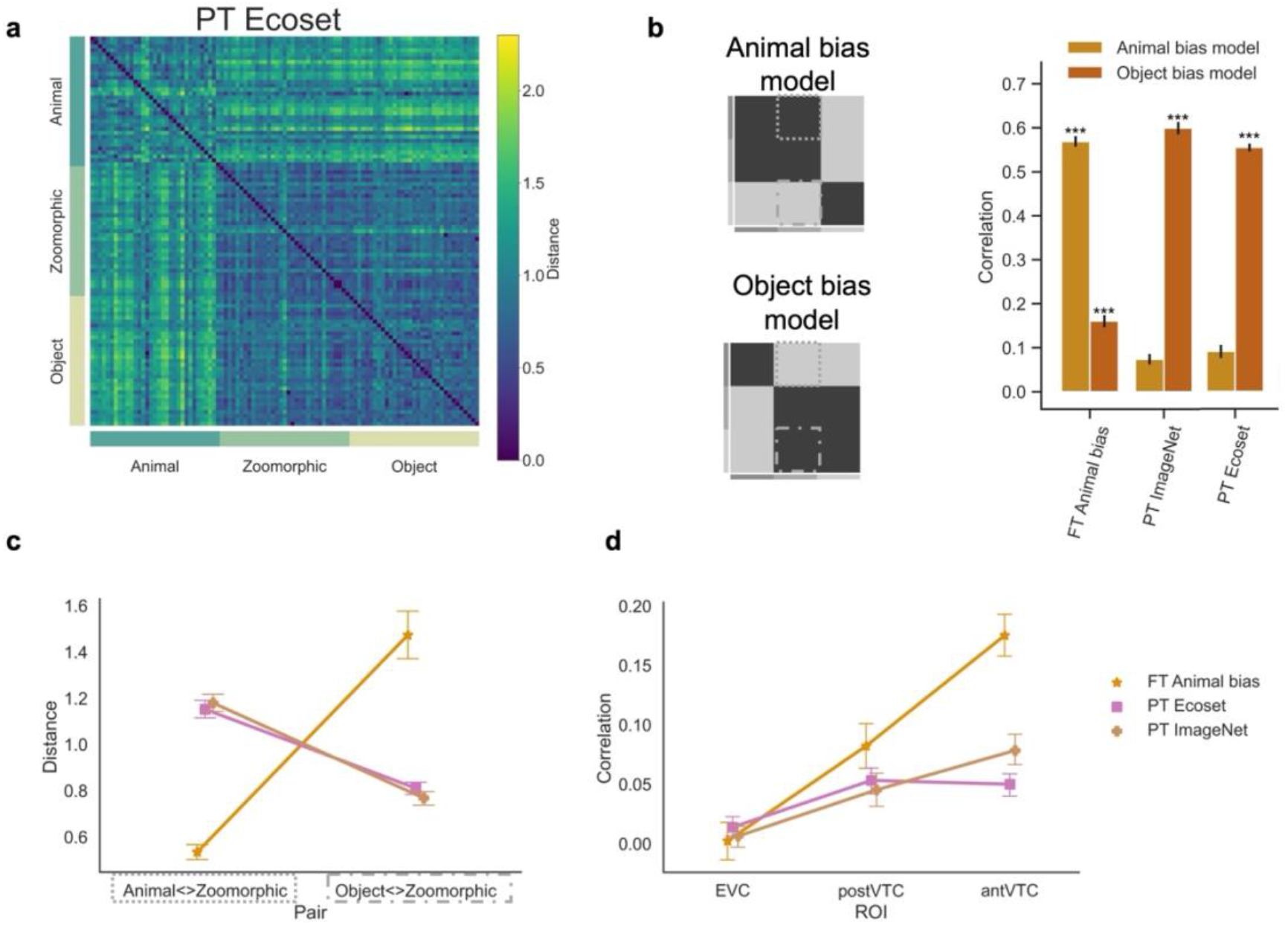
Alternative hypothesis three: training with ecological categories. **a** The RDM of the pretrained Ecoset network included in the analyses is displayed. PT = pretrained AlexNet, FT = fine-tuned AlexNet. **b** Graphical illustration of the independent Animal and Object bias model. The graph represents the correlation for each DNN with each bias model. Significant values (i.e., *** p < .0001, ** p < .001, * p < .01), were computed with permutation tests (10,000 randomizations), and error bars indicate standard error computed via bootstrapping. **c** Mean distance score of each image in the contrast Animal<>Zoomorphic and Object<>Zoomorphic and this for each DNN. **d** Graph with the individual correlations between the neural data in the three regions of interest (i.e., EVC, posterior- and anterior VTC) and the DNN data resulting from the different training regimes. All error bars represent the standard error.

In quantitative analyses, we computed the correlation with the conceptual models (Fig. 5b) and observed positive correlations with both models but much more convincingly with the Object bias model (r = .56) than with the Animal bias model (r = .09, p_paired_ < .001). Second, the Animal<>Zoomorphic and Object<>Zoomorphic distances (Fig. 5c) of the stimuli were investigated in an ANOVA with interacting factors Distances (Animal<>Zoomorphic; Object<>Zoomorphic) and Network (PT Ecoset; PT ImageNet; FT Animal bias). As for the previous results sections, a significant interaction effect was found (F(2,64) = 118.16, p < .001) and further analyzed in post-hoc analyses. Results showed that animals and zoomorphic representations were farther apart in PT Ecoset DNN (M = 1.15) relative to FT Animal bias DNN (M = 0.53, t(32) = -16.43, p_corr_ < .001), but similar to PT ImageNet (M = 1.18, t(32) = -0.95, p_corr_ = .404). Further, the distance between zoomorphic and real objects was much smaller in PT Ecoset DNN (M = 0.81) compared to FT Animal bias DNN (M = 1.47, t(32) = -4.00, p_corr_ < .001) but did not differ from PT ImageNet (M = 0.77, t(32) = 2.46, p_corr_ = .025). These results reveal that zoomorphic objects were represented more like real objects than like animals in the Ecoset pretrained DNN, in line with the Object bias observed in the standard pretrained ImageNet DNN.

Third, the correlation with the neural data (Fig. 5d) was investigated and similar results to PT ImageNet DNN were observed. A repeated measures ANOVA with factors ROI (EVC; postVTC; antVTC) x Network (PT Ecoset; PT ImageNet; FT Animal bias) revealed a significant interaction effect (F(4,60) = 15.00, p_corr_ < .001), which was further analyzed post-hoc. In anterior VTC, PT Ecoset DNN (r = 0.05) correlated significantly less compared to FT Animal bias (r = 0.18, t(15) = -7.41, p_corr_ < .001) and to PT ImageNet DNN (r = 0.08, t(15) = - 4.03, p_corr_ = .003). In posterior VTC, no significant differences were found between PT Ecoset DNN (r = 0.05) and FT Animal bias DNN (r = 0.08, t(15) =-1.77, p_corr_ = .17) nor with PT ImageNet DNN (r = 0.05, t(15) = 1.47, p_corr_ = .24). To conclude, pretrained Ecoset DNN was not able to capture more of the representations of zoomorphic objects in ant VTC than standard pretrained DNNs and performed even worse. Together, these results confirm the superiority of the Animal bias model in capturing high-level representations in visual cortex.

## Discussion

In this work, we followed a computational approach to understand the tendency of the human visual system to represent zoomorphic objects as real animals. We investigated whether and under which conditions a training protocol in DNNs can shift the representation of zoomorphic objects towards animals rather than objects. Training a DNN to explicitly induce categorize zoomorphic objects as animals increased the similarity to the Animal bias found in the brain. Obtaining such a computational Animal bias model allowed us to test a variety of other training protocols motivated by properties of human object perception. Only the explicit Animal bias training showed a shift towards an Animal bias model and by far the most convergence with anterior VTC representations. These are not just small quantitative differences, but findings that point to a representation of a different nature. While there were reasons to be very optimistic about the abilities of ImageNet-trained networks to perform object classification tasks and the convergence of these networks with perceptual and neural object representations (36,37), and researchers were rightfully impressed by further improvements due to adjusted training protocols e.g. by addressing a texture bias (34) and including more ecologically valid stimulus sets(35), all these previous developments resulted in networks with tendencies opposite to human representations when it comes to the representation of zoomorphic objects. We find that explicit supervised training to classify zoomorphic objects as animals is a crucial addition, as such providing an unprecedented understanding of the human perception of zoomorphic objects.

Our findings leave open several possibilities in terms of the exact learning mechanisms that might be involved. Effects found through supervised learning can also result from unsupervised protocols. For example, Konkle and Alvarez(38) showed that in fully self-supervised models trained on individual images, categories and hierarchical features alike those in the ventral visual stream emerged as observed in category-supervised models. The same might be true for human development, or, even more likely, a mixture of supervised and unsupervised learning might occur. The high frequency with which human infants encounter zoomorphic objects, in many cases even before they have seen the real animals, gives plenty opportunity for supervised as well as unsupervised learning mechanisms to mold the development of an Animal bias in the human visual system. Furthermore, what we model as a developmental training effect might for the real human visual system have emerged across an evolutionary time scale. A bias to, when in doubt, interpret an object as animate, might have proven important for survival in order to not miss potential predators or prey. Our computational findings are equally consistent with this evolutionary hypothesis. Further work in infants and other primate species could shed light on this issue. For now, we tend to consider the developmental hypothesis as the most likely option given the massive effects of learning on category selectivity for e.g., faces (39).

This study included four alternative hypotheses for the Animal bias based on prominent differences between the human visual system and the DNNs. Tackling the differences provided insight in whether they played a role in obtaining an Animal bias in zoomorphic objects. Two out of four human-DNN differences were related to the strong selectivity for animacy and for faces and bodies in the human brain (16), distinctions not included in the standard ImageNet training protocol. Fine-tuning pretrained networks to these distinctions did not lead to zoomorphic objects to be represented closer to animals than to objects.

The third alternative revolved around the well-documented shape – texture bias difference between DNNs and human observers(34). We hypothesized that the general shape bias in human observers might show a relationship with the Animal bias found in the human visual system for zoomorphic stimuli. However, the main training regime that has been shown previously to partially remediate the texture bias in neural networks did not induce an Animal bias for zoomorphic objects in our study. While this was an important alternative to exclude given the attention given to the texture bias, it is with hindsight maybe not surprising that an Animal bias transcends the presence of a texture or shape bias because zoomorphism can involve shape as well as texture patterns.

The last alternative training protocol included a training on a more ecologically valid stimulus set, Ecoset. This set includes distinctions that are important for human perception, such as man versus woman versus child, rather than the many tens of bird species and dog breeds that characterize ImageNet. The previously introduced neural network pretrained on this Ecoset stimulus set proved a better model for human neural representations in earlier work(35). This was not the case in the current study, to the contrary this model correlated slightly less with the neural data in anterior VTC than the older PT ImageNet network. Most importantly, the neural network trained on Ecoset totally failed to show an Animal bias with zoomorphic objects and represented such objects as objects and not as animals.

In sum, the tendency to represent zoomorphic objects as animals is a particularly prominent feature of human object processing that is only observed in neural networks when the similarity between zoomorphic objects and animals is emphasized explicitly by the training regime. Our computational analyses show that this human animal bias stands apart as a separate phenomenon that does not piggyback on other known causes of human-unique aspects of visual object cognition.

## Methods

### Stimuli

This study comprised different stimulus sets, collected from different sources (Fig. 1a). Here we only describe the stimulus sets compiled specifically for this study (Fig. 1b), in the section on DNN training regimes we will rely upon additional stimulus sets available in the literature. There were five main stimulus categories: animals, objects, zoomorphic objects, faces, and bodies. For the categories of animals, objects and human faces, the Caltech-256 stimulus set (40)was used, which has 257 categories, with on average 119 images per category. For each category (i.e., animal, object, human face), we looked for the Caltech categories that fell under this category label. In a next step, all the different images from these Caltech categories were stored as one category (e.g., animals). As a result, our main categories did not contain subcategories (e.g., different animals), but one big set of all images (e.g., animals) together. The faces subcategory animal faces were a random sample of the Animal Faces-HQ stimulus set (41). The remaining categories (zoomorphic objects and bodies) were the result of a google search with the use of the Image Downloader Chrome Extension to download the images in bulk, which were later manually evaluated whether they matched our search criteria.

For training and fine-tuning (FT) the neural networks, we created different stimulus sets that involve a subset of categories or combination of categories (Fig 6a). First, the FT Animal bias set consisted of two categories, namely objects and a combination of fifty-fifty animal and zoomorphic images put together as one category. Second, the FT Object bias contained the same images as FT Animal bias, but organized here as a pure animal category and a mixed category with objects and zoomorphic objects. Third, FT Animacy had two categories, namely animals and objects, which did not contain any zoomorphic images. Last, FT Face-body-object consisted of three categories: faces (animal + human), bodies (animal + human), and objects. In order to train and validate the networks of interest, all stimulus sets created here, were further split into a training (‘train’) and validation (‘val’) folder (Fig 1c).

Another and last collection of stimuli was created to test our DNNs (Fig 1c). Importantly, none of these images were part of training or validating the networks, and were only used to evaluate our hypotheses (see Results). This collection consisted of three categories: animals, zoomorphic objects, and objects. Each category contained 33 images, of which nine were identical to the stimuli used in the study by Bracci and colleagues (17). This subset with nine items was used to correlate the DNN data with the neural data from Bracci and colleagues (17).

### DNN data

Convolutional deep neural networks have been trained in object classification tasks, in which they reach very high levels of proficiency even with only a ‘small’ number of layers (37). AlexNet, which won the ILSVCR2012 with the complexity of ‘only’ eight layers, is the first exemplar of this generation of networks and since its conception also the one that has been used most in comparisons with human behavior and neuroscience (37). AlexNet is due to its limited complexity, and good object recognition an ideal candidate for this study. Of particular interest for our purposes are the later/deeper layers, also called fully connected layers, as these are well suited for object classification (42–46)and have been shown to correspond best to representations found in human ventral visual cortex (23). In Supplementary Fig. 1, we show that our findings do not critically depend on this architecture or layer, given that we replicated the main findings with another architecture (VGG) and for layer FC7 (Supplementary Fig. 2).

### DNN training regimes

We included multiple training regimes, resulting in multiple DNNs that share the same architecture but have a different training history. We must make a distinction between pretrained networks and fine-tuned networks. Pretrained networks were trained from scratch on a very large stimulus set, in this case ImageNet, Stylized ImageNet, or Ecoset. ImageNet consists of _∼_1.2 million images, divided over 1000 image categories, of which ∼40% animals and ∼60% objects. Stylized ImageNet was created by Geirhos and colleagues (34) to limited to texture bias present in DNNs. In short, all images were transformed for the local texture cues to be useless in object categorization. With Ecoset, Mehrer and colleagues (35), constructed a stimulus set (i.e., 565 categories) in which the distribution of concepts is more ecologically valid (i.e., frequent in language and concrete) for human daily life, which in turn yielded slightly better models of human vision. The pretrained DNNs were not trained inhouse, but rather collected from the internet: pretrained AlexNet on ImageNet via PyTorch, Stylized ImageNet through the Geirhos GitHub page (https://github.com/rgeirhos/texture-vs-shape), and PT Ecoset through the Kietzmann lab website (https://www.kietzmannlab.org/Ecoset/). The baseline pretrained network is trained on ImageNet (PT ImageNet), whereas Stylized ImageNet (PT Stylized ImageNet) was included to tackle the texture bias common in DNNs, and Ecoset (PT Ecoset) was the ecologically valid alternative that better matched models of human vision.

The baseline PT ImageNet served as the starting point for our fine-tuned networks, in which we further trained the PT ImageNet with our different training stimulus sets (Fig. 1b): animals + zoomorphic <> objects (FT Animal bias), animals <> zoomorphic + objects (FT Object bias), animals <> objects (FT Animacy), and faces <> bodies <> objects (FT Face-body-object). This fine tuning is also known in the literature as transfer learning, and it makes use of a well-established trained network to further train the network on a similar, but smaller stimulus set. In this approach, the weights of most of the layers are not changed. This approach is beneficial when using a small but similar stimulus set as the early layers are well trained for basic image features and do not need to change as a result of the new training.

The current study is interested in ‘deeper’ brain regions along the visual processing pathway, such as ventral occipitotemporal cortex. Other advantages are that this approach is less time consuming and prevents overfitting of the network. Given the small size of the stimulus set, all layers are set frozen except the penultimate fully connected layer (=FC7) and the last fully connected output layer (=FC8). FC8 is now characterized by the number of categories in the stimulus set (i.e., 2 or 3) rather than 1000 in pretrained (Stylized) ImageNet networks, and 565 in pretrained Ecoset. Fine-tuning was initialized with a learning rate of 0.01, which was gradually multiplied by a factor 0.1 in case the validation loss did not significantly improve over five consecutive epochs with a minimum learning rate of 0.00001. The training applied a batch size of 20 over 500 epochs, with a Stochastic Gradient Descent optimization algorithm, Cross Entropy Loss function, momentum of 0.9, and data is reshuffled at every epoch, resulting in a top-1 validation accuracy ranging between 90.9% and 99.5% (see Table 1).

**Table 1.** Overview of the top-1 validation accuracies (%) for AlexNet for the different fine-tuned networks.

| Network | Validation accuracy (%) |
| --- | --- |
| FT Object bias | 90.9 |
| FT Animal bias | 93.2 |
| FT Animacy | 93.1 |
| FT Face-body-object | 99.5 |

### DNN representational dissimilarity matrices

The test set with 99 images (i.e., 33 untrained images in each category: animal, zoomorphic, object) was loaded into the differently trained DNNs (i.e., PT ImageNet, PT Stylized ImageNet, PT Ecoset, FT Animal bias, FT Object bias, FT Animacy, and FT Face-body-object) to obtain the image representations from the output layer. These representations include the responses of all nodes in a layer, and thus varied in size depending on the number of nodes/categories in the network output layer (i.e. respectively 1000, 1000, 565, 2, 2, 2, and 3 nodes). For each network, a representational dissimilarity matrix (RDM) was constructed by calculating the distance score for each image pair (1 – Pearson R). In a next step, these RDMs were normalized by dividing each score by the mean of the entire RDM.

A subset of the RDM, only containing the stimuli from Bracci and colleagues (17), was used when correlating with the neural data available for that subset (17). Additionally, during these analyses, these subset RDMs were limited to the upper triangle with exclusion of the diagonal.

The data obtained through DNNs are different from human data, as there is only one RDM per network (i.e., no individual data available like in human research). We applied non-parametric statistics for analyses that only entail DNN data. Permutation statistics across stimuli are used for correlation analyses, whereas more advanced non-parametric statistics for ANOVA were included to test multi-factorial models that take into account variability across stimuli. More specifically, through the ARTool software (47,48), the dependent variable (i.e., distance score) was transformed (i.e., align and rank) to perform a non-parametric factorial ANOVA and later post-hoc analyses, which are corrected for multiple comparisons (i.e., BH FDR).

### Description of the data from Bracci et al. (2019)

Bracci and colleagues (17) investigated the perceptual phenomenon ‘perceiving animacy’ in the human behavior and brain, and in DNNs. For this, they made use of independent models of animacy (hereinafter referred to as “Object bias model”) and appearance (hereinafter referred to as “Animal bias model”), which are in a sense RDMs containing zeros and ones for ideal cases where zero stands for 100% like each other and one for 0% similar. The Object bias model (Fig 2b) expects great similarity along the category diagonal and between objects and zoomorphic objects, whereas the Animal bias model (Fig. 2b) expects great similarity between animals and zoomorphic objects next to the diagonal similarity. The RDMs coming from the behavioral, neural and DNN experiment were correlated with these independent model RDMs.

At the DNN level, two pretrained DNNs (i.e., VGG-19 and GoogLeNet) were evaluated with the Bracci test subset. Here a strong correlation was found with the Object bias model, whereas there was no sign of any correlation with the Animal bias model. In the neuroimaging part, subjects performed two different tasks in the scanner: an animacy task (“does this image depict a living animal?”; inducing an Object bias) and an appearance task (“does this image look like an animal?”; inducing an Animal bias). Bracci et al. (17) obtained RDM data from three regions of interest: early visual cortex (EVC), posterior (post-) and anterior (ant-) ventral temporal cortex (VTC). While EVC served as a control region of low-level features, VTC has shown its importance in object representation and categorization (49). Results showed correlations in VTC with both models, but unexpectedly significantly stronger for the Animal bias model. Furthermore, when looking at post and ant VTC separately, results are even more pronounced in ant VTC where there is no longer a correlation with the Object bias model.

In our study, the new DNN data was correlated with the existing neural data. The neural RDMs consisted of an average across both tasks given that there was no effect of task in any of the ROIs.

## Data and software

All DNN work, except for Ecoset, and analyses were done in PyTorch (v1.9.0) and Torchvision (v0.10.0) through Spyder (Python 3.8). For Ecoset, we used Code Ocean (https://codeocean.com/capsule/9570390/tree/v1) to extract the image vectors from the pretrained AlexNet on Ecoset. For conducting non-parametric factorial ANOVA and corresponding post-hoc analyses we used ARTool (47,48). The data used for final statistics are made available through the Open Science Framework at https://osf.io/7xcg9/?view_only=5df7e38a5f91411eaee71b22aa34eda7.

## Acknowledgements

This work was supported by the EOS Grant HUMVISCAT to HO), the KU Leuven Grant (C14/21/047 to HO) and the Fund for Scientific Research – Flanders (11G6219N to SD) (https://www.fwo.be/). The funders had no role in study design, data collection and analysis, decision to publish, or preparation of the manuscript.

## Author Contributions

Most work was done by the first author, supervised by HO. SD collected the stimuli, trained and tested all the network instances, analyzed the data, and created the figures. SD and HO jointly designed the training regimes, interpreted the results, and wrote the manuscript together with SB.

