## Supplementary Fig. for "A computational understanding of zoomorphic perception in the human brain"

### Supporting Information

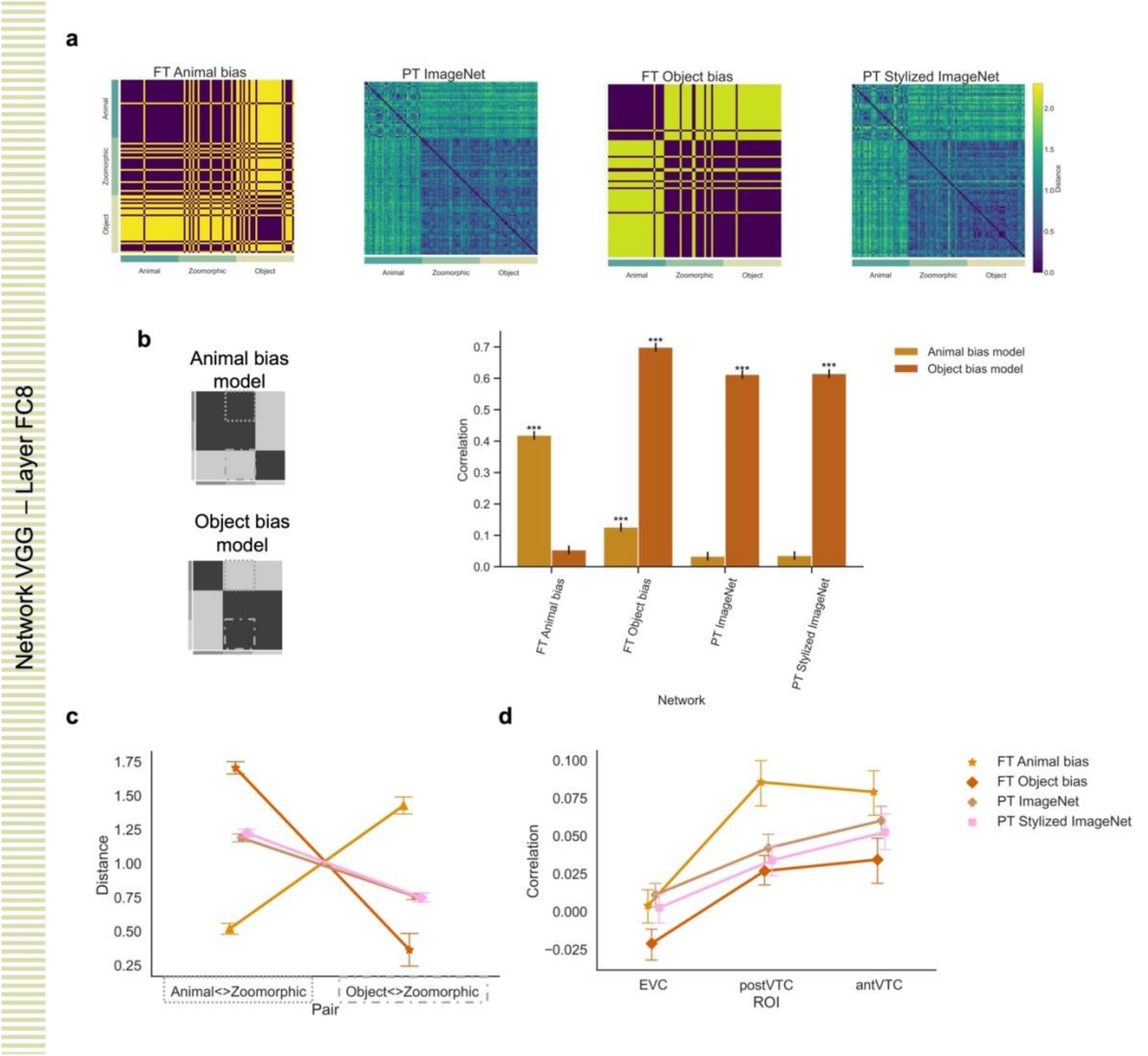


**Supplementary Figure 1. Overview of main findings in VGG layer FC8. a** The RDMs of four networks included in the analyses are displayed. PT = pretrained VGG, FT = fine-tuned VGG. **b** Graphical display of the independent Animal bias model and Object bias model. **c** The graph represents the correlation for each DNN with each bias model. Significant values (i.e., *** p < .0001, ** p < .001, * p < .01), were computed with permutation tests (10,000 randomizations), and error bars indicate standard error computed via bootstrapping. **d** Mean distance score of each image in the contrast Animal<>Zoomorphic and Object<>Zoomorphic and this for each DNN, showing the same effect as for AlexNet so that only for FT Animal Bias we see a strongly upward curve (Animal<>zoomorphic smaller than

Object<>Zoomorphic). **e** Graph with the individual correlations between the neural data in the three regions of interest (i.e., EVC, posterior- and anterior VTC) and the DNN data resulting from the different training regimes. For VGG we also found, as before for AlexNet, that FT Animal Bias has the strongest correlation with the neural data, although in case of VGG this effect is most prominent for posterior VTC. All error bars represent the standard error. VGG-19 is used for all DNNs, except for PT Stylized ImageNet, which was available in VGG-16.

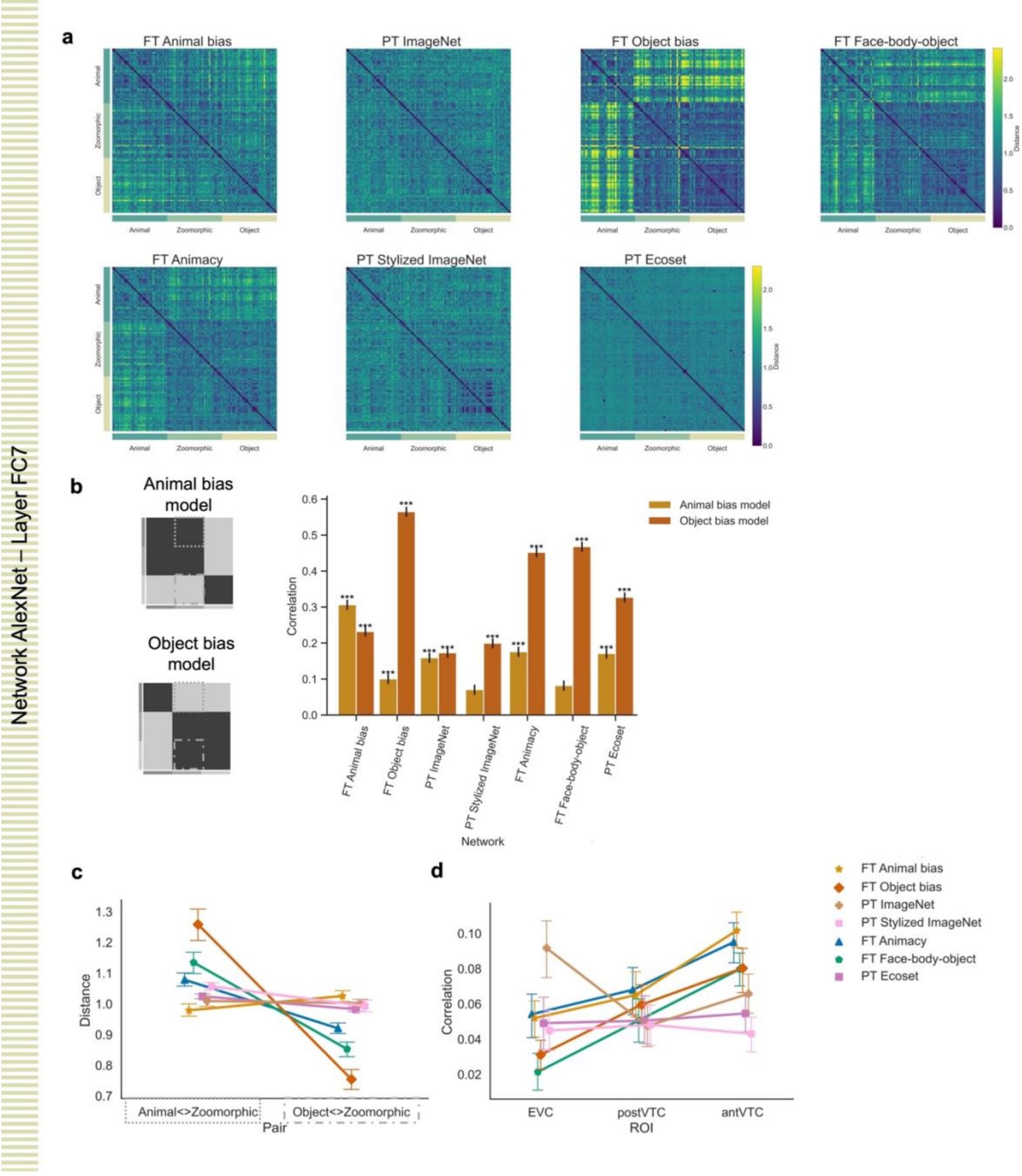

**Supplementary Figure 2. Overview of main and alternative findings in AlexNet layer FC7. a** The RDMs of three main networks included in the analyses are displayed. PT = pretrained AlexNet, FT = fine-tuned AlexNet. **b** Graphical display of the independent Animal bias model and Object bias model. **c** The graph represents the correlation for all tested DNNs with each bias model. Significant values (i.e., *** $p < .0001$, ** $p < .001$, * $p < .01$), were computed with permutation tests (10,000 randomizations), and error bars indicate standard error computed via bootstrapping. **d** Mean distance score of each image in the contrast Animal<>Zoomorphic and Object<>Zoomorphic and this for each DNN. Also in FC7 we find that FT Animal Bias tends to have the most upward going curve (Animal<>zoomorphic smaller than Object<>Zoomorphic), but the difference with the other networks is much smaller than for FC8. **e** Graph with the individual correlations between the neural data in the three regions of interest (i.e., EVC, posterior- and anterior VTC) and the DNN data resulting from the different training regimes. Also in FC7 the correlation with VTC representations tends to be the highest for FT Animal Bias, but again the effects are small compared to findings in FC8. The relative effect size in different layers might depend upon meta-parameters such as learning rate and how many layers are fixed during fine-tuning, yet it is reassuring that effects go in similar directions in different layers.

All error bars represent the standard error.
